# The *Arabidopsis thaliana* NIP2;1 Lactic Acid Channel promotes Plant Survival Under Low Oxygen Stress

**DOI:** 10.1101/2020.08.03.234641

**Authors:** Zachary Beamer, Pratyush Routray, Won-Gyu Choi, Margaret K. Spangler, Ansul Lokdarshi, Daniel M. Roberts

## Abstract

Under anaerobic stress *Arabidopsis thaliana* induces the expression of a collection of core hypoxia genes that encode proteins associated with an adaptive response. Included in these core hypoxia genes is *NIP2;1*, which encodes a member of the “Nodulin-like Intrinsic Protein” (NIP) subgroup of the aquaporin superfamily of membrane channel proteins. Under normal growth, *NIP2;1* expression is limited to the “anoxia core” region of the root stele, but shows substantial induction in response to low oxygen stress (as high as 1000-fold by 2-4 hr of hypoxia challenge), and accumulates in all root tissues. During hypoxia, *NIP2;1-GFP*, accumulates on the cell surface by 2 hr and then is distributed between the cell surface and internal membranes during sustained hypoxia, and remains elevated in root tissues through 4 hrs of reoxygenation recovery. T-DNA insertional mutant *nip2;1* plants show elevation of lactic acid within root tissues, and a reduced efflux of lactic acid and acidification of the external medium. Together with previous biochemical evidence demonstrating that NIP2;1 has lactic acid permease activity, the present work supports the hypothesis that the protein facilitates the release of cellular lactate to the rhizosphere to prevent lactic acid toxicity. In support of this, *nip2;1* plants show poorer survival to argon-induced hypoxia stress. *Nip2;1* mutant plants also show elevated expression of ethanolic fermentation transcripts, as well as reduced expression the lactate metabolic enzyme GOX3, suggesting that the altered efflux of lactate through NIP2;1 regulates other pyruvate and lactate metabolism pathways.

**One-sentence Summary:** The NIP2;1 lactic acid permease is necessary for an optimum response to low oxygen stress through the release of lactate from roots during hypoxia stress.

## Introduction

As obligate aerobes, land plants require a continuous supply of oxygen to support energy demand. Oxygen deprivation stress from flooding or poor soil aeration depresses cellular respiration leading to an energy crisis that triggers a variety of genetic, metabolic, and developmental adaptation strategies (Voesenek and Bailey-Serres, 2013, 2015). In Arabidopsis, low oxygen stress triggers the preferential transcription and translation of a collection of “core” hypoxia-induced genes that encode glycolytic/fermentation enzymes, other metabolic proteins, and various signal transduction proteins and transcription factors involved in the adaptive response (Mustroph et al., 2009; Mustroph et al., 2010). Among the hypoxia-induced core response transcripts is an aquaporin-like membrane channel protein, Nodulin Intrinsic Protein 2;1 (Choi and Roberts, 2007).

Nodulin 26-like” Intrinsic Proteins (NIPs) are a terrestrial plant-specific subfamily of the aquaporin-superfamily that show structural and functional homology to the soybean root nodule-specific protein nodulin 26 (Roberts and Routray, 2017). NIPs possess the canonical aquaporin “hourglass” fold, but have diverged as multifunctional transporters of a wide variety of substrates including glycerol, ammonia, and a variety of essential as well as boric acid, silicic acid, and toxic metalloid hydroxides (Roberts and Routray, 2017; Pommerrenig et al., 2020). Based on structural modelling, NIPs can be segregated into three “pore families” that have conserved amino acids within their aromatic-arginine (ar/R) selectivity filters (Roberts and Routray, 2017). NIP2;1, along with NIP1;1, NIP1;2, NIP3;1, NIP4;1 and NIP4;2, belongs to the Arabidopsis NIP I subgroup (Johanson et al., 2001; Routray and Roberts, 2017). NIP I proteins have an ar/R amino acid composition similar to nodulin 26 (Wallace and Roberts, 2004), and show a transport selectivity for water, glycerol, and ammonia as natural substrates (Roberts and Routray, 2017), as well as the ability to be permeated by toxic metalloids (e.g., arsenite and antimonite) (Kamiya and Fujiwara, 2009; Kamiya et al., 2009; Xu et al., 2015). Biochemical analysis of NIP2;1 in Xenopus oocytes showed that it lacks the typical NIP I transport properties and is impermeable to classical NIP substrates (Choi and Roberts, 2007). Instead, NIP2;1 shows pH-dependent transport of lactic acid (Choi and Roberts, 2007).

Lactic acid is the end product of lactate fermentation, one the fermentative pathways employed by plants to sustain energy production under oxygen limiting conditions and other stress conditions in which respiration is repressed (Drew, 1997; Gibbs and Greenway, 2003). Accumulation of lactic acid increases the acid load in the cytosol, and is among the factors that contribute to the cellular acidification observed during low oxygen stress (Davies et al., 1974; Roberts et al., 1984; Felle, 2005). Hypoxia-induced fermentation is accompanied by the acquisition of the ability of plant roots to efflux lactic acid to the rhizosphere (Xia and Saglio, 1992; Rivoal and Hanson, 1993; Xia and Roberts, 1994; Dolferus et al., 2008). In maize root tips, this lactic acid efflux mechanism is correlated with reduced susceptibility to low oxygen stress from acclimation, it is proposed to be an adaptive mechanism to reduce cytosolic acidification or other toxic effects of cellular lactic acid/lactate accumulation (Xia and Saglio, 1992; Xia and Roberts, 1994).

The previous finding that NIP2;1 is selectively permeable to lactic acid upon expression in oocytes, together with its identification as a core-hypoxia gene product, has led to the hypothesis that it mediates the efflux and/or compartmentation of lactic acid during the Arabidopsis response to low oxygen stress (Choi and Roberts, 2007). However, this has yet to be investigated in planta. The objective of this research was to test this hypothetical function of NIP2;1 during the Arabidopsis hypoxia response. In this study, it shown that a T-DNA insertional *nip2;1* mutant exhibits poor tolerance of low oxygen stress compared to wild-type. Further, it is shown that the efflux of lactate from hypoxia-stressed roots to the media from the roots requires NIP2;1 expression. NIP2;1 localization was observed to vary between internal membranes and the plasma membrane during the course of hypoxia and recovery, consistent with previous observations suggesting dual localization of the protein. Lastly, evidence is presented that the transcripts encoding fermentation and pyruvate/alanine metabolism enzymes are altered in *nip2;1* mutant plants compared to wild-type plants. Overall, the data support a role for NIP2;1 as a lactic acid efflux channel, and suggest that this activity could help control the expression of enzymes that metabolize pyruvate during the Arabidopsis hypoxia response.

## Results

### *NIP2;1* expression is induced early during hypoxia primarily in root tissue

Hypoxia treatment of two week old Arabidopsis seedlings resulted in a rapid increase in *NIP2;1* expression in root tissues with Q-PCR analysis revealing a >1000-fold increase in *NIP2;1* transcript levels within two hours after the onset of anaerobiosis (Fig. 1A). *NIP2;1* transcript levels then showed a sharp decline by 12 hr but still remained over 100-fold elevated relative to normoxic controls before leveling off at 24 hr (Fig. 1A). While *NIP2;1* is predominantly a root transcript, hypoxia also induced *NIP2;1* expression in shoot tissues, but with a lower overall expression (40-fold relative to basal levels) compared to roots (Fig. 1A inset). In addition, the time course of accumulation in shoots was delayed when compared to root tissues and expression did not peak until 12 hr treatment.

**Figure 1.**
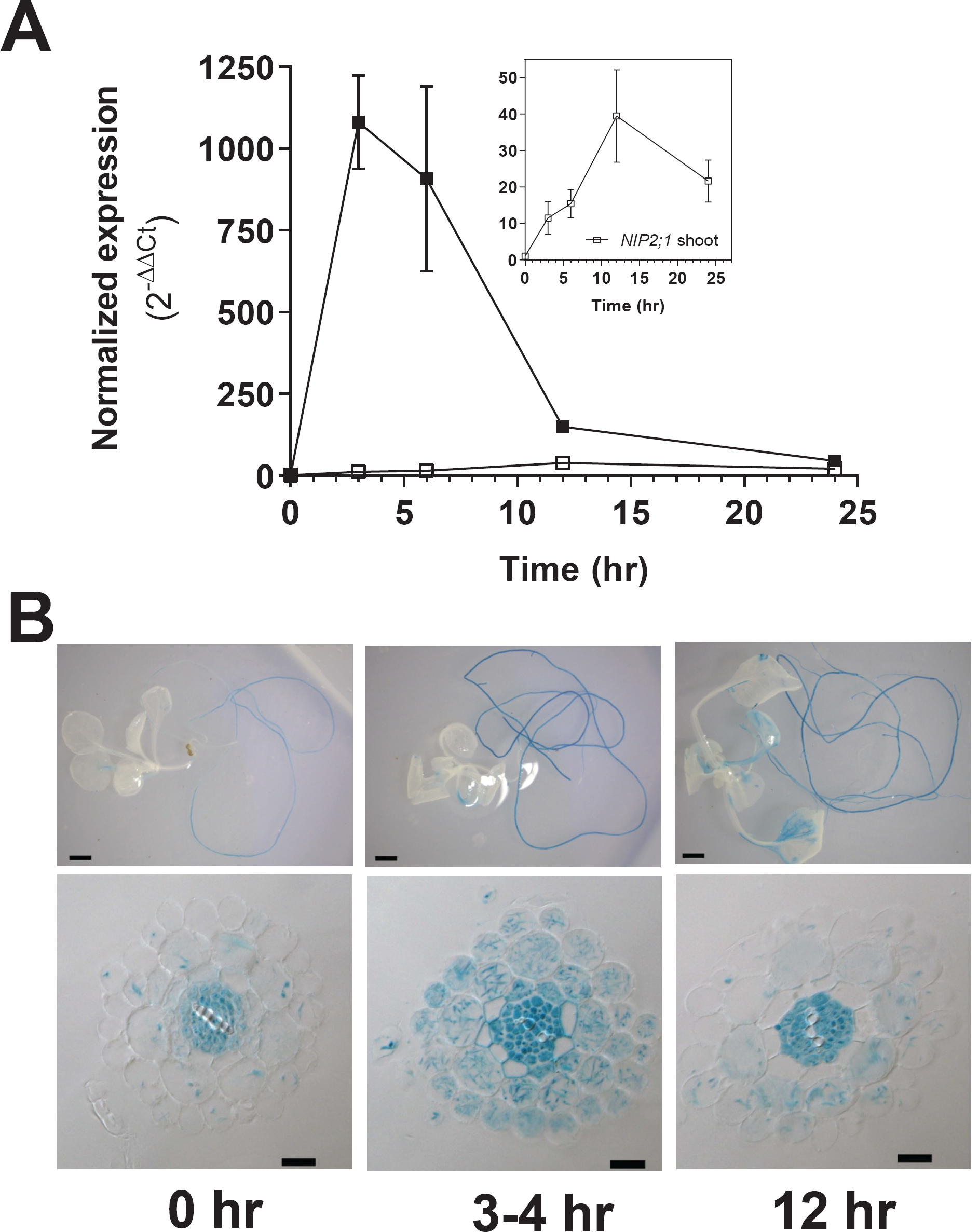
*NIP2;1* expression in Arabidopsis seedlings in response to oxygen deficit. **A**. Quantitative real-time RT-PCR (Q-PCR) analysis of *NIP2;1* transcripts in root (filled squares) and shoot (open squares) tissues during a hypoxia time course of two week old Arabidopsis seedlings. The ΔCt value of AtNIP2;1 obtained from 0 hr of shoot sample was used as the calibrator for ΔΔCt calculations. The graph in the insert shows a rescaled plot of *NIP2;1* expression in shoot tissue. Error bars indicate the SD of three biological replicates. **B**. GUS staining analysis of two-week old *NIP2;1pro::GUS* Arabidopsis seedlings subjected to the oxygen deprivation conditions as in panel A. Top panel are representative whole seedlings while the bottom panel shows root cross-sections. Scale bars are 1.0 mm for the top panel and 20 μm in the bottom panel.

Similar patterns of *AtNIP2;1* expression were observed with two week old *AtNIP2;1 promoter::GUS* fusion plants subjected to the same oxygen deprivation regime (Fig 1B). Analysis of the cellular localization of GUS staining under normoxic conditions showed that expression is principally limited to roots with little staining detected in shoot tissues (Fig. 1B). Cross-sections of unstressed (normoxic) roots showed that GUS staining was principally observed in the cells of the stele (pericycle, phloem and procambium,) with little or no GUS signal apparent in endodermal, cortical and epidermal cells (Supplemental Fig. 1, Fig. 1B). At 4 hr after the induction of hypoxia, root tissues showed an increase in intensity of the GUS staining in the stele as well as the appearance of the GUS signal in the cortex, epidermis and root hairs (Fig. 1B). Similar to the Q-PCR result, this staining peaked at 4 hr post hypoxic treatment and then decreased by 12 hr post treatment (Fig. 1B), although the expression at these later time points was still much higher than the basal expression in the unstressed roots (Fig. 1B). In comparison to roots, increases in GUS expression in shoots were less acute and appeared more slowly and the expression was mainly restricted to the vascular tissues of leaves (Fig. 1C).

### *NIP2;1* enhances plant survival under low oxygen conditions

Core hypoxia-response gene loss-of-function mutants generally result in reduced survival or increased sensitivity to low oxygen stress (Ismond et al., 2003; Kursteiner et al., 2003; Licausi et al., 2010; Giuntoli et al., 2014; Sorenson and Bailey-Serres, 2014; Lokdarshi et al., 2016). To investigate whether *NIP2;1* is necessary for hypoxia stress survival, a T-DNA insertion mutant (WiscDsLox233237_22K; referred to here as *nip2;1)* line was studied. The *nip2;1* line contains a T-DNA insertion within the promoter region (−30) between a cluster of Anaerobic Response Elements and the transcriptional start site (Fig. 2A). Consistent with the position of this insertion, hypoxia treatment (4 hr) of *nip2;1* plants resulted in poor expression of *NIP2;1* compared to wild type plants (Fig. 2B). Assay of NIP2;1 protein levels by Western blot showed no detectable NIP2;1 protein in root tissues of *nip2;1* mutant plants after 6 hours of hypoxia compared to wild type plants (Fig. 2C).

**Figure 2.**
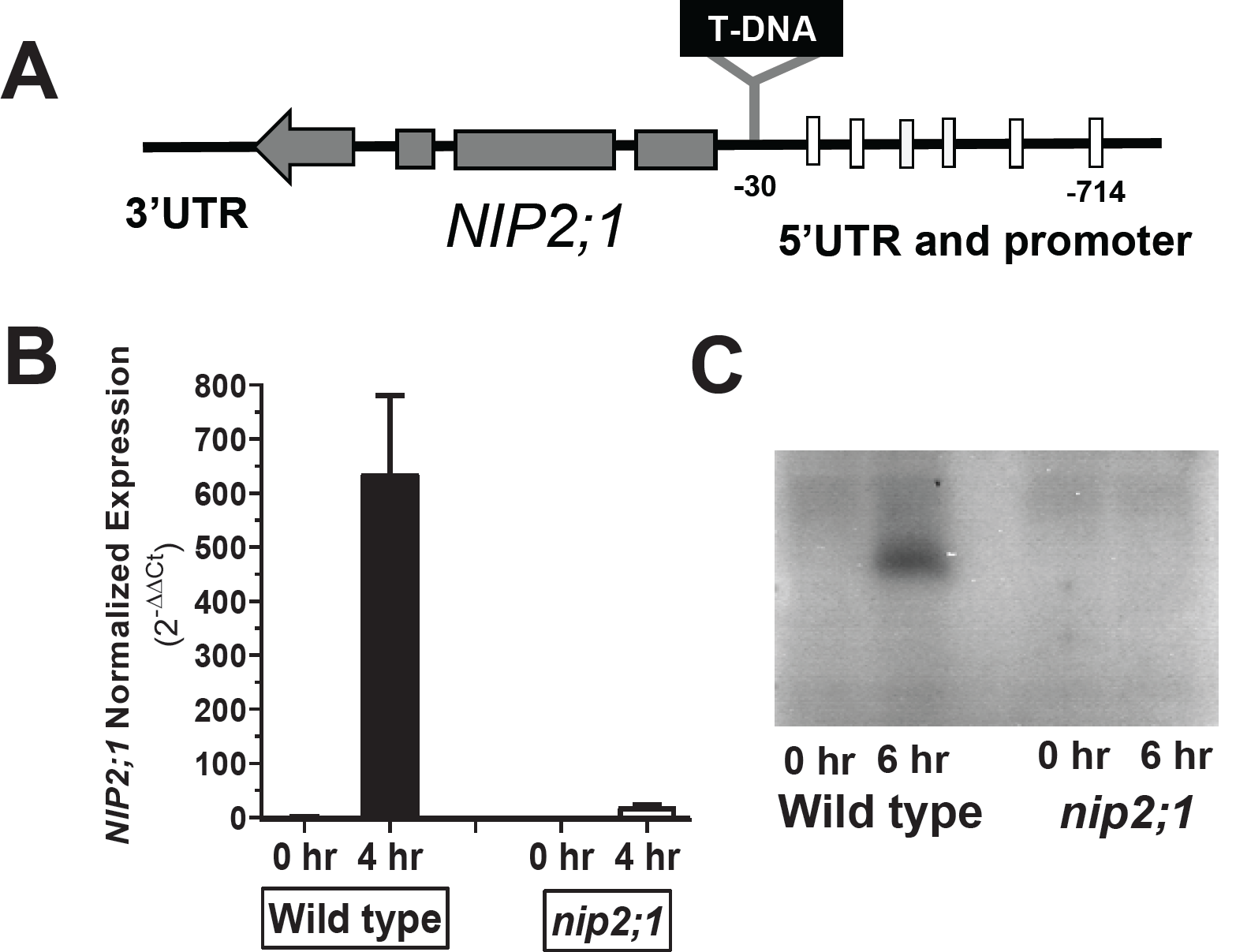
Characterization of *nip2;1* T-DNA insertional mutant seedlings. **A**. schematic diagram showing the T-DNA insertion in *nip2;1* mutant line. The boxes in the upstream portion of the gene indicate Anaerobic Response Elements. **B**. Q-PCR results for *NIP2;1* expression in the roots of 7d-old wild type and *nip2;1* during hypoxia treatment. The ΔCt value of *NIP2;1* expression in normoxic wild type roots was used as a calibrator for relative expression. Error bars show the SD of 3 biological replicates. **C**. Root extracts (10 μg protein/lane) were analyzed by Western blot with site-directed NIP2;1 antibodies. **0 hr**, normoxic control; **6 hr**, 6 hr hypoxia-treated plants.

To determine the effects of low oxygen stress on *nip2;1* plants, their growth and survival under normoxic and hypoxic conditions were compared (Fig. 3). While wild type and *nip2;1* plants showed little difference in growth under normoxic conditions (Supplemental Fig 2), *nip2;1* plants showed higher sensitivity to hypoxia treatment (Fig. 3). After exposure to argon gas-induced hypoxia, followed by transfer back to normoxic conditions, *nip2;1* seedlings exhibited a higher incidence of leaf chlorosis within 24 hours after the hypoxia treatment (Fig. 3A). Comparison of the overall survival frequency of wild type and *nip2;1* seedlings showed that the mutant exhibited significantly poorer survival to hypoxic stress (Fig. 3C).

**Figure 3.**
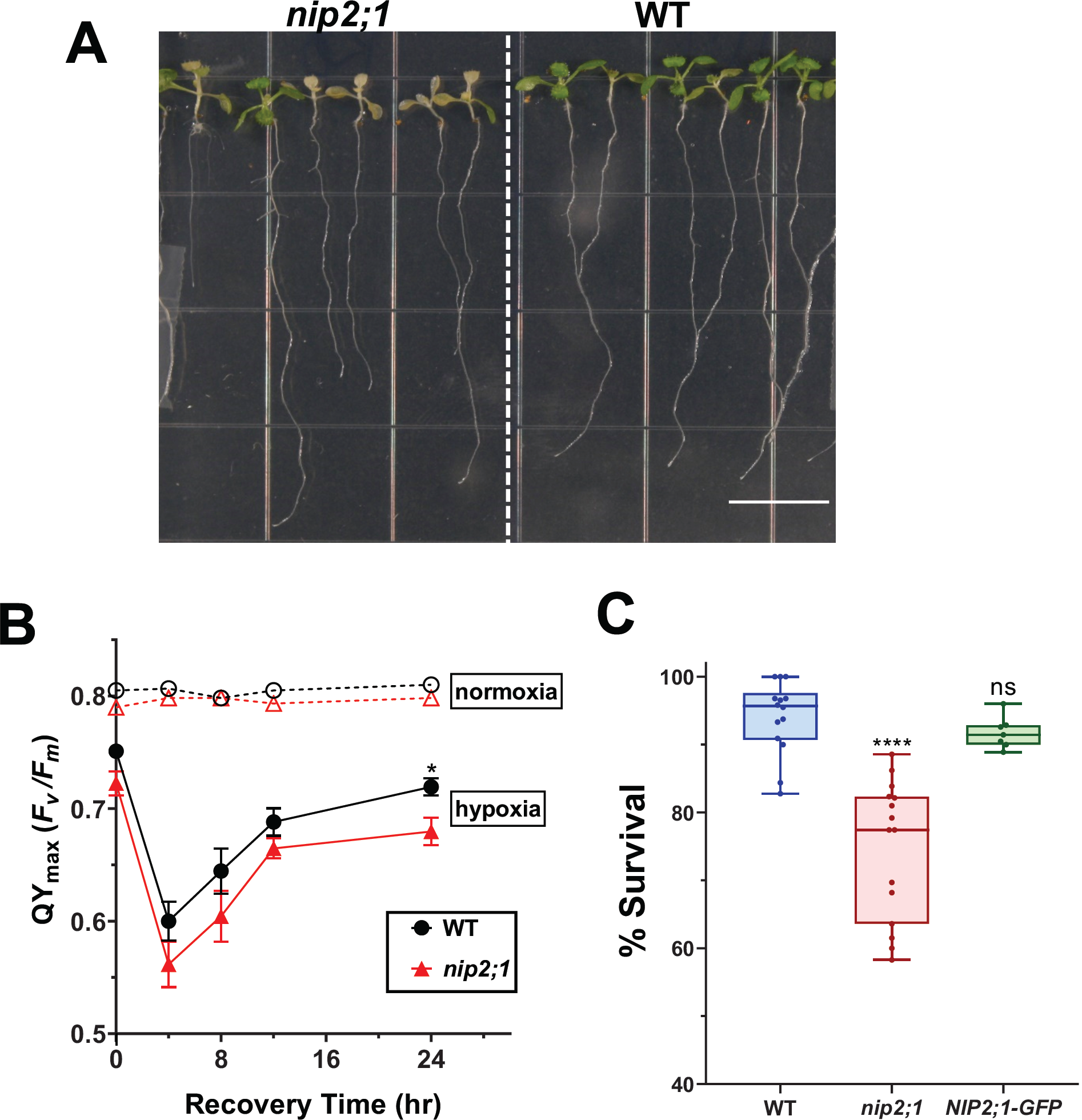
Effects of oxygen deprivation on the survival of *nip2;1-1* T-DNA insertional mutant seedlings. **A**. Seven-day-old, vertically grown seedlings corresponding to the wild-type (WT) and *nip2;1* T-DNA insertional mutant (*nip2;1*) were subjected to 8 hrs of argon gas treatment and were allowed to recover under normal oxygen conditions for 72 hours. Scale bar = 1 cm **B**. PSII maximal quantum yield, QY_max_ (F_v_/F_m_) was calculated from chlorophyll fluorescence analysis of 16hr light, 8hr dark grown 7-days-old WT and *nip2;1* mutant seedlings treated with 8 hr argon in dark (hypoxia) or air control (normoxia). Error bars represent std. error of mean of five biological replicates. Statistical significance was assessed by student t-test analysis to the wild type survival value. (* represents p<0.05) **C**. Histogram showing the survival of seven day old WT, *nip2;1*, and *NIP2;1-GFP* complementation seedlings to 8 hr argon treatment represented as a box and whisker plot of % seedling survival. Each data point represents one biological replicate with the median value indicated in each box. Statistical significance was assessed by student t-test analysis to the wild type survival value. (**** represents p<0.0001; ns, not significant).

To confirm that the increased sensitivity of *nip2;1* plants to hypoxia is result of the loss of *NIP2;1* gene, complementation lines containing a *NIP2;1pro::NIP2;1-GFP* transgene in the *nip2;1* background were analyzed (Fig. 3C and Supplemental Fig. 2). The complementation lines showed enhanced tolerance to hypoxia challenge compared to *nip2;1* plants, (Supplemental Figure 2), and were not statistically different to wild type plants with respect to survival frequency (Fig. 3C).

The sensitivity of the *nip2;1* mutant to hypoxia was further assessed by measuring the chlorophyll fluorescence properties and calculating the maximum potential quantum efficiency (Q_max_ or F_v_/F_m_) of photosystem (PS) II of *nip2;1* and WT seedlings under normal and low oxygen stress conditions. Chlorophyll fluorescence measurements is a common technique used to assess the photosynthetic efficiency of PS II which is an index of the susceptibility of plants to different environmental stressors (Murchie and Lawson, 2013). Under normoxic conditions, WT and *nip2;1* seedlings were not significantly different with F_v_/F_m_ values that fell within the optimum range (0.78 - 0.8, Murchie and Lawson, 2013). This suggests that the lack of *NIP2;1* expression in the mutant line does not exhibit any detectable adverse effect on this parameter under standard growth conditions. However, upon hypoxia treatment both WT and *nip2;1* showed a steep reduction in photosynthetic efficiency with the F_v_/F_m_ ratio declining to below 0.6 within four hours of the recovery period (Fig. 3B). At all time points, the F_v_/F_m_ ratio is lower for *nip2;1* than wild type plants. At later time points the quantum efficiency of PS II starts to recover (Fig. 3C), but the *nip2;1* F_v_/F_m_ ratio remains significantly lower than wild type after 24 hours of recovery.

Taken together, the *nip2;1* phenotype data indicate that *NIP2;1* is hypoxia core response protein that participates in the hypoxia adaptation response, and that reduction in expression of *NIP2;1* lowers the ability of Arabidopsis to survive this stress.

### NIP2;1 is expressed on the plasma membrane as well as on internal membranes during hypoxia and reoxygenation recovery

To investigate the dynamics of protein expression and the subcellular localization of NIP2;1, the *NIP2;1pro::NIP2;1-GFP* complementation lines were analyzed. Similar to wild type *NIP2;1*, Q-PCR analysis shows that the *NIP2;1-GFP* transgene transcript is acutely induced by hypoxia, with a peak at 4 hr followed by a decline (Fig. 4C). As has been documented with other core hypoxia transcripts(Branco-Price et al., 2008), reoxygenation results in a rapid decline of the transcript to basal levels. Analysis of NIP2;1-GFP protein accumulation during hypoxia and re-oxygenation was done by using epifluorescence microscopy and Western blot analysis at different time points of hypoxia stress and reoxygenation recovery (Fig. 4A and B). The GFP signal first appeared within two hours of the onset of hypoxia (Fig. 4A and B), and increased as hypoxia proceeded. Return to normal oxygen conditions resulted in a substantial increase in the fluorescent intensity at 30 min (Fig. 4A) with Western blot protein levels peaking at 2 hr reoxygenation (Fig. 4B and C). While transcript levels decline rapidly to non-detectable levels (Fig. 4C), the fluorescent intensity and Western blot analyses indicate the persistence of the NIP2;1 protein for hours after return to normal oxygen conditions (Fig. 4A and B).

**Figure 4.**
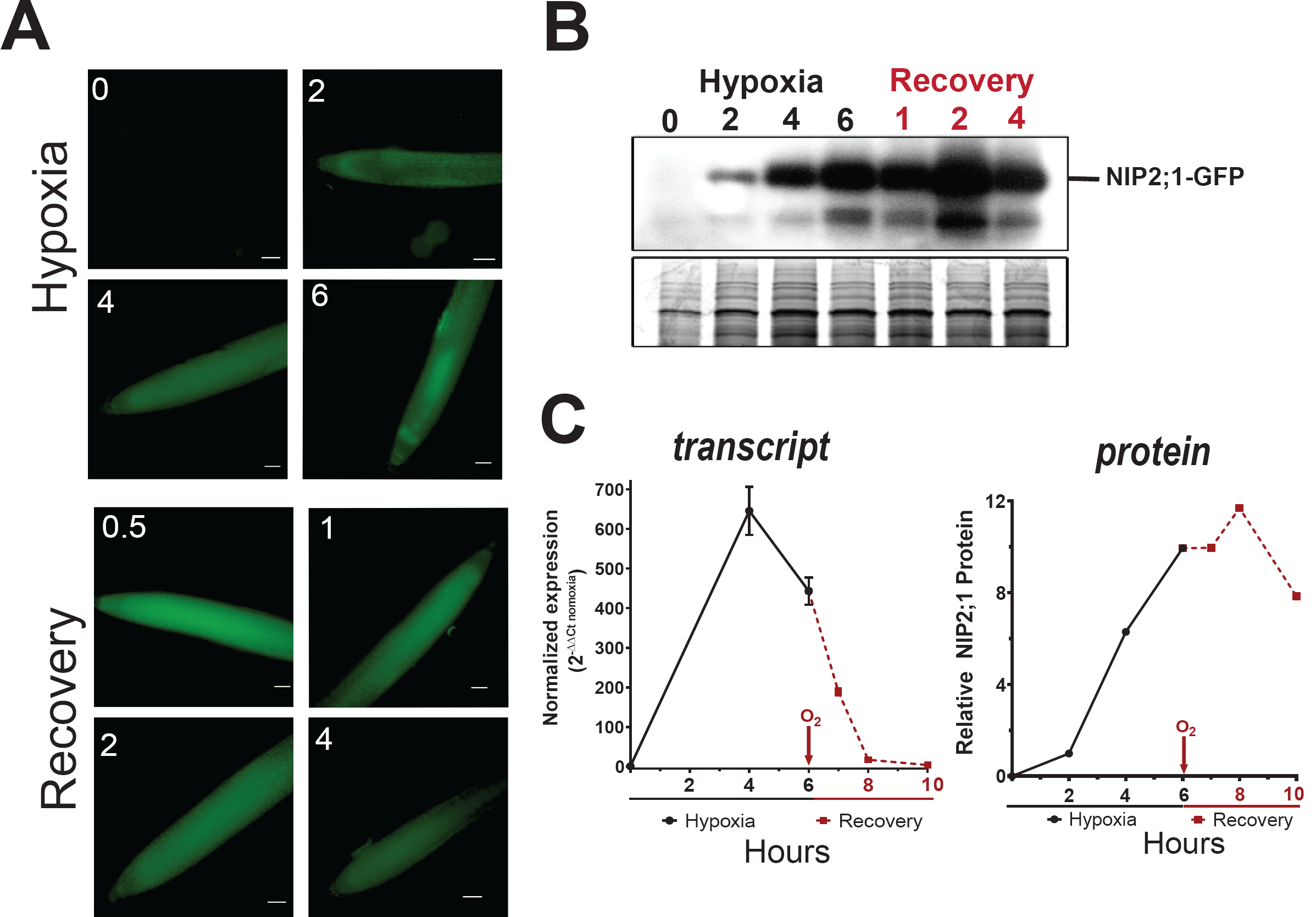
NIP2;1-GFP expression in the roots of *NIP2;1:GFP* plants during hypoxia and reoxy-genation. Ten day old *NIP2;1-GFP* plants were subjected to an argon-induced hypoxia time course, with oxygen resupplied at hour 6. **A**. Representative epifluorescent images of the primary root of *NIP2;1:GFP* plants at the indicated times of hypoxia treatment or reoxygenation recovery. Scale bars = 50 μm. **B**. Anti-GFP Western blot showing NIP2;1-GFP protein accumulation (upper panel). Bottom panel, Coomassie blue stained loading control gel. **C**. Comparsion of root *NIP2;1-GFP* transcript levels (normalized to 0 hr) by Q-PCR (*left*) and NIP2;1-GFP protein based on densitometry of the Western blot iin panel B.

Different NIP proteins show varied subcellular localization, ranging from polarized plasma membrane localization for the root boric acid permease NIP5;1(Wang et al., 2017) to specific localization on subcellular organelles such as the soybean symbiosome membrane protein, nodulin 26 (Weaver et al., 1991). Previous work with NIP2;1 expression in heterologous systems with strong, constitutive non-native promoters suggested localization to either the plasma membrane (Choi and Roberts, 2007) or internal membranes (Mizutani et al., 2006). To investigate its subcellular localization under native conditions during hypoxia and reoxygenation, confocal microscopy of the roots of *NIP2;1pro::NIP2;1-GFP* seedlings was done (Fig. 5 and Supplemental Fig. 3).

**Figure 5.**
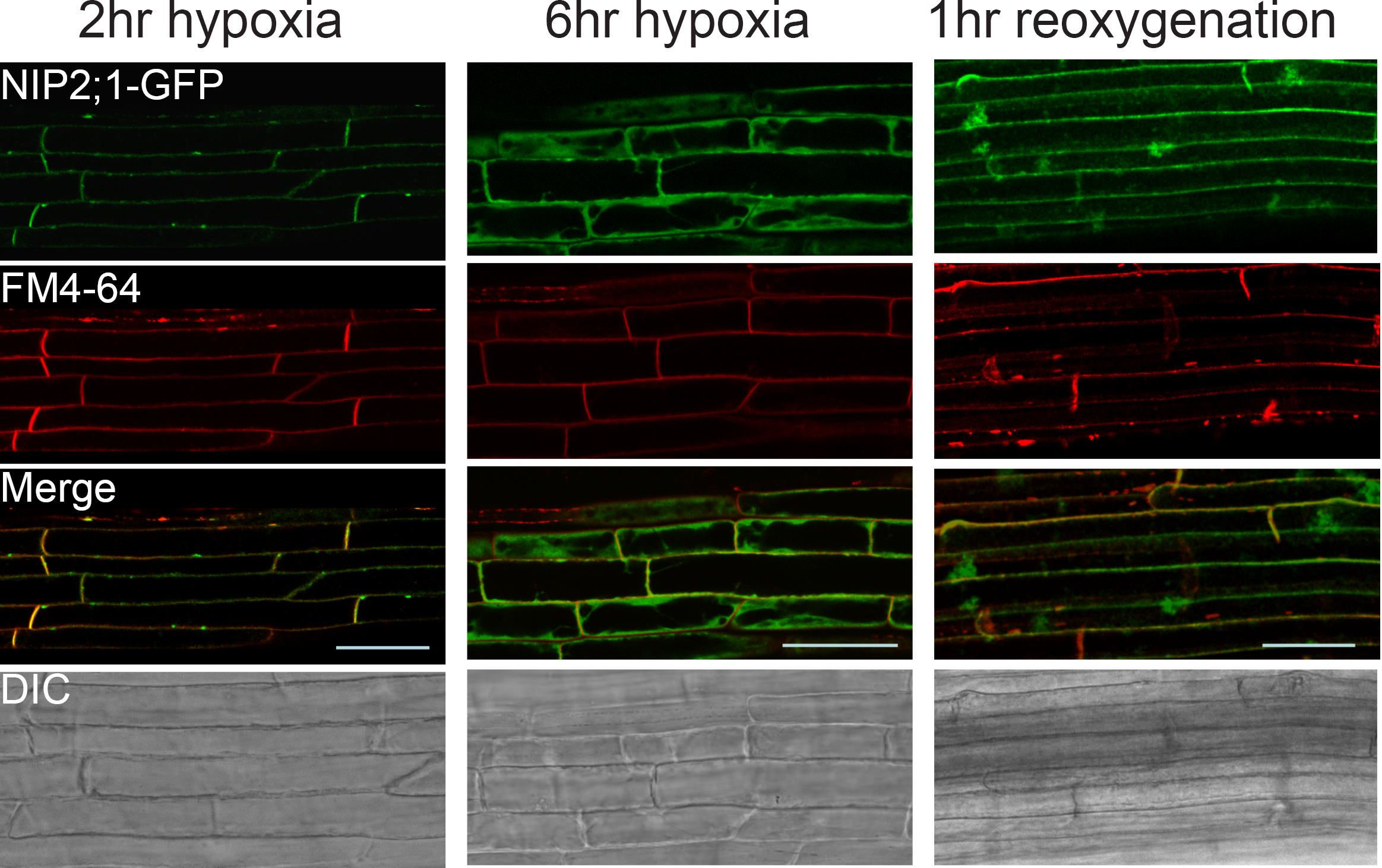
Subcellular localization of NIP2;1-GFP in the roots of hypoxia challenged 7-d-old *NIP2;1-GFP* complementation lines. Seven-day-old vertically grown *NIP2;1-GFP* seedlings were subjected to anaerobic stress by root submergence under argon gas treatment followed by return to aerobic conditions at 6 hr. The appearance of NIP2;1 GFP was monitored at the times indicated by confocal fluorescence microscopy of the root elongation zone. FM4-64 staining was used to visualize the plasma membrane. DIC, differential interference contrast images. Bars = 50 μm.

Analysis of NIP2;1-GFP at 2 hr after the onset of hypoxia revealed accumulation within the elongation zone of the root with significant co-localization at the plasma membrane with the marker FM4-64 (Supplemental Fig. 3, Fig. 5), although some staining within internal structures is also apparent (Supplemental Fig. 3B). As hypoxia proceeds and more protein accumulates, the majority of NIP2;1-GFP signal appears to accumulate internally (Fig. 5, 6 hr time point). Following 1 hr of reoxygenation, the localization of NIP2;1 on the surface appears to increase, although internal localization is still apparent (Fig. 5).

**Figure 6.**
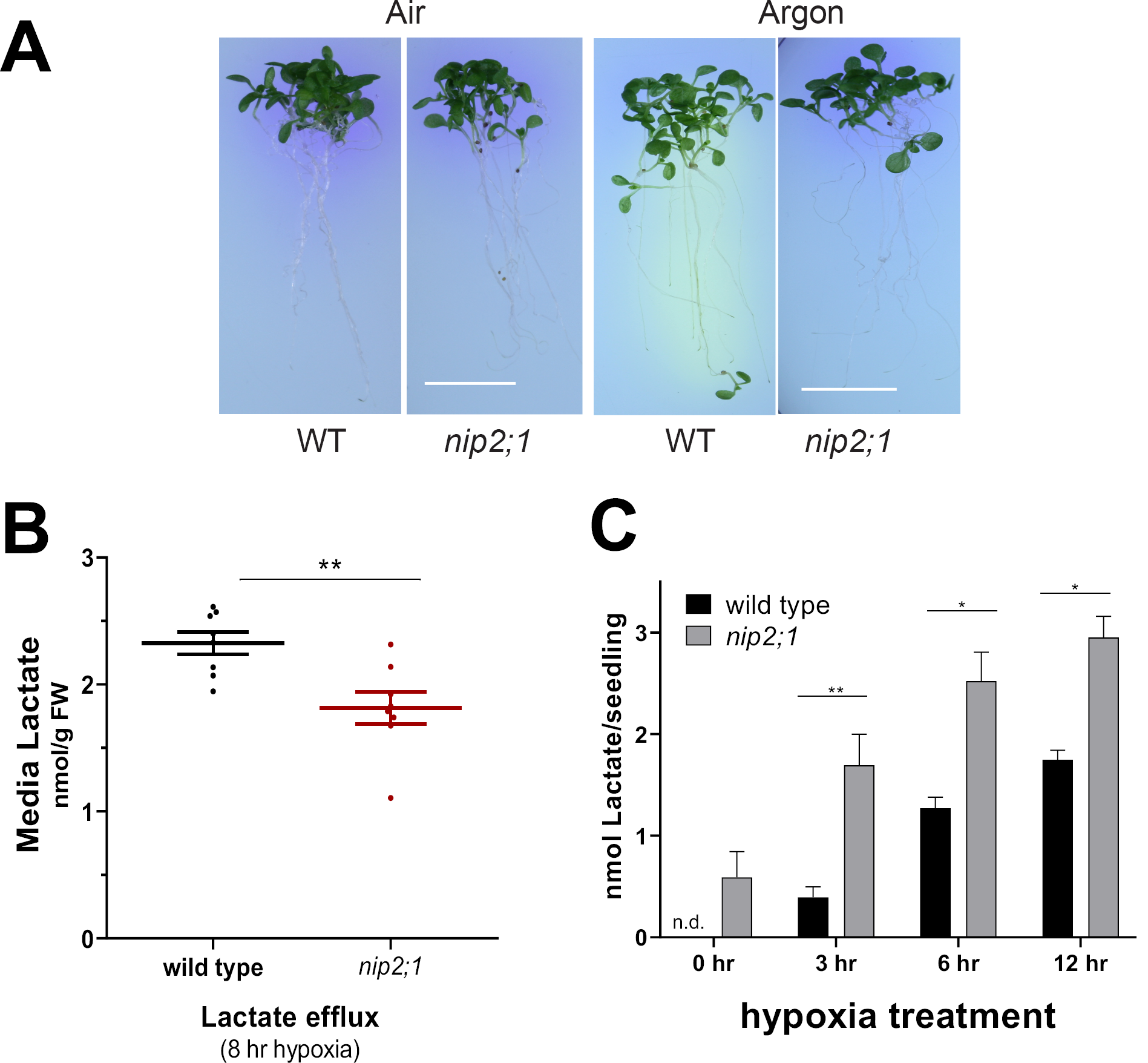
Media acidification and lactic acid efflux in hypoxia-challenged *nip2;1* and wild type plants. **A**. Ten day-old *nip2;1* and wild-type (WT) seedlings were transferred to pH indicator plates containing bromocresol purple, and were subjected to 8 h treatment of hypoxia induced by argon gas **(Argon**). **Air** indicates a normoxic control. A color change from purple to yellow indicates a decrease in the pH of the environment. Scale bar = 1 cm. **B**. 10-d-old *nip2;1* and wild-type seedlings were submerged in water and subjected to argon gas treatment for 8 hr and the lactate concentration in the media was measured. Data are represented as a scatter plot of eight biological replicates with the horizontal line representing the mean and error bars showing the SE. **C**. Accumulation of lactate in seedlings based on enzymatic analysis during hypoxia induced by bubbling nitrogen gas. Values are the average of three biological replicates at each timepoint, with the error bars showing the SEM. Asterisks indicate statistically significant differences based on a paired Student’s t-test analysis (panel B) or One way ANOVA analysis (panel C).

### NIP2;1 participates in lactate efflux and media acidification during hypoxia

Based on the specificity of NIP2;1 as a lactic acid permease based on biochemical analyses (Choi and Roberts, 2007), its localization in part to the surface of root cells, and the observation that hypoxia triggers release of lactate from roots into the external media (Xia and Saglio, 1992; Dolferus et al., 2008; Engqvist et al., 2015), it is hypothesized that NIP2;1 may participate in the excretion of lactate during the low oxygen stress response. To test this hypothesis, the pH and concentration of lactic acid within the media of wild-type and *nip2;1* seedlings challenged with hypoxia were examined. To compare the hypoxia-induced acidification of the external medium, 10 d-old wild-type and *nip2;1* seedlings were subjected to argon gas treatment on media containing the pH sensitive dye, bromocresol purple (Fig. 6A). Upon transfer from normoxic to hypoxic conditions, wild type plants show significant acidification of the media surrounding the roots, while *nip2;1* plants show no difference in bromocresol purple staining between normoxic and hypoxic conditions (Fig 6A).

In follow up analyses, the lactate levels released into the media by the roots of hypoxia-challenged wild type and *nip2;1* seedlings were compared. Wild type roots excrete significantly higher amounts of lactate into the environment compared with the roots of *nip2;1* plants (Fig 6B). The levels of lactate that accumulate within wild type and *nip2;1* roots tissues under hypoxic conditions were also compared (Fig. 6C). During a hypoxia time course, lactate levels increase in both wild type and *nip2;1* roots, but at all time points, including normoxia, *nip2;1* plants accumulate significantly higher levels of this metabolite (Fig. 6C). The results, combined with previous functional studies (Choi and Roberts, 2007), suggest that NIP2;1 is an endogenous lactic acid transporting channel protein that participates in lactate transport, homeostasis, and efflux during low oxygen stress in Arabidopsis roots, and that this activity is necessary to release lactate produced during anaerobic fermentation within the root.

### The loss of *NIP2;1* function affects the expression of pyruvate and lactate metabolic enzymes

Anaerobic metabolism of pyruvate during oxygen limitation in plants occurs through three conserved pathways: lactic acid fermentation, ethanolic fermentation and alanine synthesis (Fig. 7). While all three pathways use pyruvate as a substrate, lactic acid and ethanolic fermentation regenerate NAD^+^, whereas alanine synthesis serves as a mechanism to store nitrogen and carbon for reoxygenation (Sato et al., 2002; Ricoult et al., 2005). Genes that encode enzymes in these pathways (such as *ADH, PDC, LDH*, and *AlaAT*) are among the “core hypoxia response” proteins that are induced in Arabidopsis roots during hypoxia (Mustroph et al., 2009; Lee et al., 2011; Mustroph et al., 2014). Conversely, L-lactate produced via LDH is proposed to be converted back to pyruvate in peroxisomes (Fig. 7A) by the root-specific glycolate oxidase 3 enzyme (Engqvist et al., 2015). Unlike other members of this enzyme family that participate in the metabolism of glycolate, GOX3 is specific for L-lactate and utilizes oxygen as an electron acceptor to oxidize lactate, producing hydrogen peroxide and pyruvate as end products (Engqvist et al., 2015). *GOX3* is not a hypoxia-induced transcript and is rather proposed to regulate the concentration of lactate in a coordinate fashion with LDH under aerobic conditions (Engqvist et al., 2015; Maurino and Engqvist, 2015).

**Figure 7.**
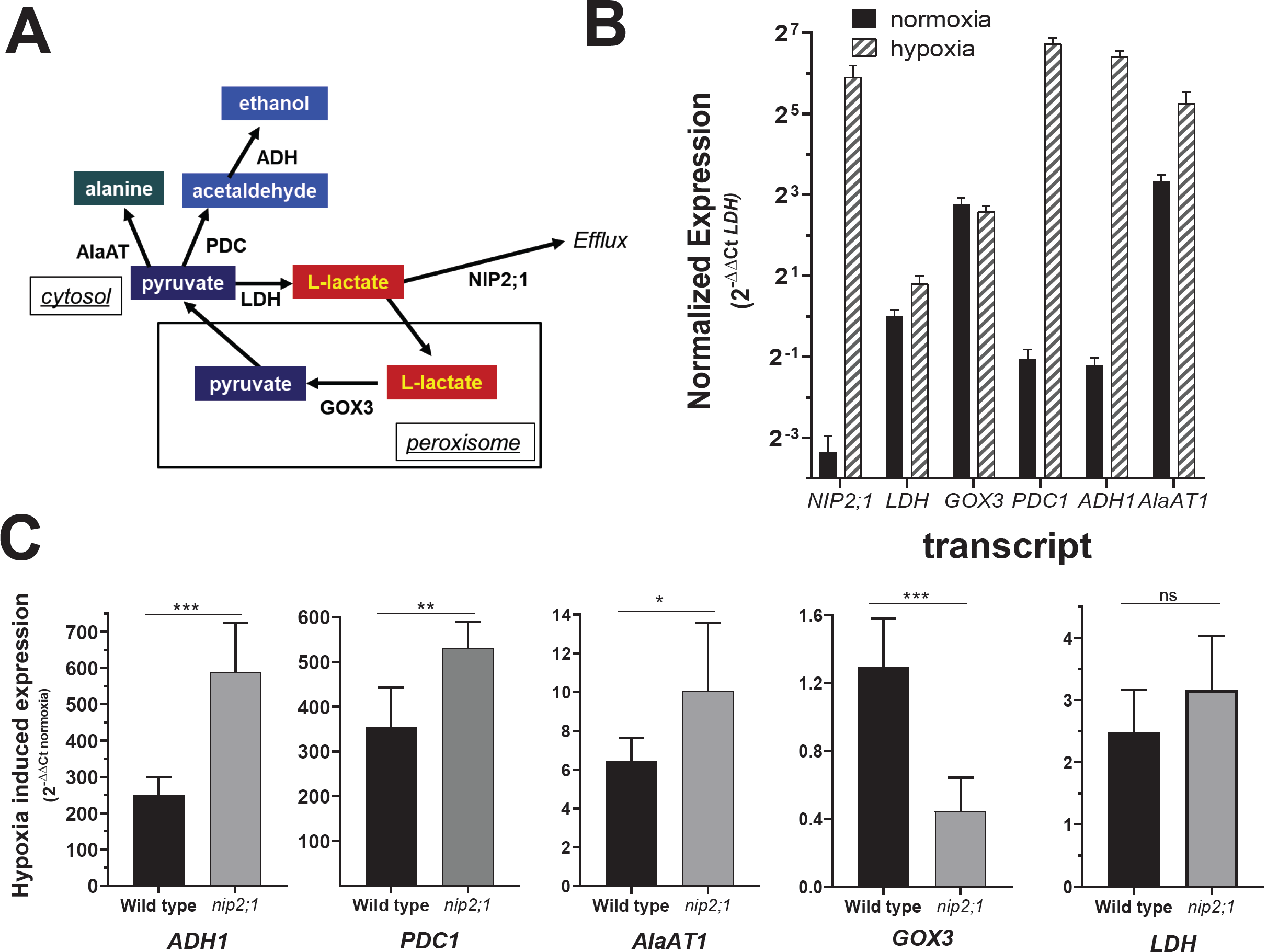
Effect of *nip2;1* mutation on transcripts of pyruvate metabolism enzymes. **A**. Scheme showing the principal pathways of pyruvate and lactate metabolism during fermentation. **B**. Q-PCR analysis of the indicated transcripts in the roots of seven day old wild-type seedlings grown under normoxic conditions (black bars) or in response to 4 hr of argon-induced hypoxia (stripped bars). The data are normalized to the transcript levels of LDH under normoxic conditions (normalized expression = 2^-ΔΔCt-LDH^). Data are the average of nine replicates with the error bars showing the SD. **C**. Comparison of the hypoxia-induced changes of selected transcripts from the roots of wild type and *nip2;1* seedlings. The data were normalized to the expression levels of the indicated transcript under normoxic conditions. Data are the average of six determinations from three biological replicates with the error bars showing the SD. Asterisks indicate statistically significant differences based on an unpaired Student’s t-test analysis.

Q-PCR analysis shows that hypoxia response transcripts encoding fermentation enzymes (*ADH1, PDC1, LDH*, and *AlaAT1*) show induction in root tissues during 4 hr of argon gas (Fig. 7B). However, closer analysis of wild type and *nip2;1* roots show different levels of selective transcripts. While *LDH* transcript levels show no statistical differences between wild type and *nip2;1* roots, *ADH1, PDC1*, and *AlaAT1* are substantially elevated in *nip2;1* compared to wild type plants (Fig. 7C). Conversely, *GOX3*, which is expressed at the same level under normal and hypoxic conditions in wild type roots, is substantially reduced in hypoxic *nip2;1* roots. Overall, the data suggest that the alterations in lactic acid homeostasis within the *nip2;1* mutant have a selective effect on the metabolism of pyruvate and lactate metabolizing enzyme expression, with transcripts encoding fermentation enzymes in ethanol and alanine producing pathways enhanced, whereas the transcript that encodes the lactic acid metabolizing enzyme GOX3 is suppressed.

## Discussion

### NIP2;1-mediated lactic acid efflux promotes Arabidopsis survival during low oxygen stress

In response to low oxygen conditions resulting from flooding or submergence stress, plants switch to anaerobic fermentation pathways to maintain glycolytic flux and energy production. In Arabidopsis, enzymes of ethanolic (ADH and PDC) and lactic acid (LDH) fermentation pathways are necessary for optimal survival to low oxygen stress(Ellis et al., 1999; Ismond et al., 2003; Kursteiner et al., 2003; Dolferus et al., 2008). The levels of lactate increase 14-fold within the root during the first two hours of hypoxia challenge (Mustroph et al., 2014) suggesting that lactic acid fermentation is induced during the initial stages of hypoxia. As lactate accumulates, hypoxia-stressed Arabidopsis plants (Dolferus et al., 2008), similar to other plant lineages (Xia and Saglio, 1992; Rivoal and Hanson, 1993; Xia and Roberts, 1994), efflux lactate from the roots to the external media/rhizosphere during low oxygen stress. Lactate efflux mechanisms in plant roots may assist in mitigating cellular acidification or other toxic effects of lactate accumulation during anaerobic stress (Xia and Roberts, 1994; Greenway and Gibbs, 2003).

To alleviate potential cellular acidification from lactate fermentation, the pathways for the efflux of lactic acid must transport either the protonated form (lactic acid) or co-transport lactate with a proton [reviewed in (Greenway and Gibbs, 2003)]. In the case of animal cells, lactate is effluxed or taken up by members of the SLC16 subgroup of the major facilitator superfamily known as Monocarboxylate Transporters or MCTs (Counillon et al., 2016; Sun et al., 2017). These proteins are symporters that co-transport lactate with H^+^ in a bidirectional fashion. They participate in the efflux of excess lactic acid during anaerobic fermentation, and also serve as an uptake mechanism for lactate from the serum for further metabolism (Sun et al., 2017). Land plants lack members of the SLC16/MCT transporter family, and the molecular identity of the transporters or channels that mediate the efflux of lactate/lactic acid produced during anaerobic fermentation have remained unclear. In this study, cellular, and genetic evidence is provided that, together with previous protein functional analyses (Choi and Roberts, 2007), indicate that the aquaporin-like NIP2;1 assists in mediating lactic acid efflux from Arabidopsis roots.

Nodulin Intrinsic Proteins are a plant-specific subgroup of the aquaporin superfamily that have diversified structurally and functionally during land plant evolution, and which have acquired solute transport activities beyond canonical aquaporin water transport (Roberts and Routray, 2017). NIP transport functions range from glycerol to ammonia to metalloid metabolites such as boric and silicic acid. However, biophysical and biochemical analysis of Arabidopsis NIP2;1 in Xenopus oocytes (Choi and Roberts, 2007) indicate that it is an outlier from other classical NIP proteins and is impermeable to water and all traditional NIP solute substrates, and instead shows specific bidirectional permeability to protonated lactic acid. Several observations in the present work provide strong support that lactic acid transport and efflux is the biological function of NIP2;1. First, the expression of NIP2;1 in response to hypoxia coincides with the appearance of lactate in the external medium; second, genetic mutation of the NIP2;1 via T-DNA insertion results in the reduction of lactate efflux from hypoxic roots into the external medium and a concomitant increase in the accumulation of lactate within root tissue; and third, mutant *nip2;1* plants show reduction in the acidification of the media surrounding hypoxic roots.

Mutant *nip2;1* plants show poorer survival to argon-induced low oxygen stress compared to wild type plants, presumably because of the over accumulation of toxic levels of lactic acid due to a reduced ability to efflux this end product from roots. As noted above, cytosolic lactate generation would increase the acid load of the cytosol that could contribute to acidosis (Davies et al., 1974; Roberts et al., 1984; Felle, 2005). Additionally, the accumulation of lactate could also contribute to reduced glycolytic flux by affecting NAD^+^ regeneration by altering the equilibrium of the LDH reaction, or potentially through product feedback inhibition mechanisms. For example, recent studies in yeast and mammals show that over accumulation of lactate leads to the production of the toxic side product 2-phospholactate catalyzed by pyruvate kinase. This side product of lactate blocks the production of fructose-2-6 bisphosphate, leading to the inhibition of the key glycolytic enzyme phosphofructokinase-1 (Collard et al., 2016). The production of similar toxic lactate metabolites side products could conceivably occur in plant tissues as well.

### NIP2;1 expression during normoxia, hypoxia stress and recovery

NIP2;1 expression is almost completely limited to root tissues with a precise pattern of transcript and protein expression during normoxic, hypoxic, and reoxygenation conditions. Under normoxic conditions, *NIP2;1* promoter activity is restricted to cells within the stele of the mature root (Supplemental Fig. 1). The cells of the stele are hypoxic even under well aerated growth conditions due to the low rate of lateral oxygen diffusion across the mature differentiated root (Armstrong et al., 1994; Gibbs and Greenway, 2003). “Anoxic cores’’ in the root stele are proposed to aid in hypoxia sensing and acclimation, potentially by the communication of low oxygen or energy signals (ethylene, metabolites, low pH, and Ca^2+^) between hypoxic and well-aerated cells (Armstrong et al., 2019). The roots of *nip2;1* mutants show increased accumulation of lactate under normoxic conditions (Fig. 6C). This suggests that LDH is active in anoxic core tissues, even under aerobic conditions, and that NIP2;1 basal expression is necessary for maintaining low lactate accumulation.

*NIP2;1* Q-PCR and GUS data show the characteristics of a core hypoxia response transcript, with acute induction of *NIP2;1* expression in roots within 1 hr of the initiation of hypoxia stress, followed by a peak and eventual decline to a reduced but elevated steady state level that is sustained during hypoxia. Interestingly, examination of the cell-specific translatome atlas based on the work of (Mustroph et al., 2009) shows that the expression *NIP2;1* during hypoxia parallels the expression of the two lactate metabolizing enzyme transcripts, *LDH* and *GOX3* (Supplemental Figure 4). All three transcripts are predominantly, if not exclusively, expressed in root tissue (Dolferus et al., 2008; Mustroph et al., 2014; Engqvist et al., 2015), and accumulate to the highest levels in the root cortex, as well as to high levels within the epidermal and vascular tissues, but are absent or poorly expression in leaf tissues. The root-predominant expression pattern of *NIP2;1, LDH1* and *GOX3* is consistent with the distinct lactate metabolic properties and responses of roots and shoots to low oxygen stress (Ellis et al., 1999; Mustroph et al., 2014). Based on the model of (Engqvist et al., 2015), these three gene products are proposed to coordinate lactate homeostasis through its production (LDH), its recovery back to pyruvate (GOX3), and the excretion of lactate from the cell when it is overproduced during low oxygen stress (NIP2;1).

Similar to other hypoxia-induced genes (Branco-Price et al., 2008), reoxygenation results in suppression of *NIP2;1* mRNA expression and a return to low basal levels within two hours of recovery. In contrast, NIP2;1 protein levels increase during early reoxygenation and remain elevated for several hours post recovery, suggesting that the activity of the protein is also required during recovery. In addition to excretion of lactic acid to the media, Arabidopsis can take up L-lactate from the media and metabolize it (Dolferus et al., 2008; Engqvist et al., 2015). Since NIP2;1 mediates the bidirectional flux of lactic acid (Choi and Roberts, 2007), it could assist in the recovery of excreted lactic acid to trigger its metabolism to pyruvate and entry into the TCA cycle as part of the replenishment of TCA cycle intermediates that takes place during post-anoxic recovery(Branco-Price et al., 2008; Tsai et al., 2014; Yeung et al., 2019).

### NIP2;1 localization during hypoxia stress and recovery

By using the complementation lines with a *NIP2;1-GFP* transgene under the native promoter, the localization of the protein during hypoxia stress and recovery was analysed. NIP2;1-GFP shows a strong colocalization with a plasma membrane marker during early hypoxia when the protein starts to appear (2 hrs). During extended hypoxia (>6 hr) the NIP2;1 GFP signal becomes stronger, and with enhanced localization within endomembranes although expression on the cell surface is still observed. During recovery, dual localization is still observed, although plasma membrane localization appears to be prevalent. These observations indicate that the NIP2;1 is distributed to both surface and internal membranes, and can explain the potential conflicting reports of different subcellular localization (plasma membrane and ER) based on transient expression in Arabidopsis protoplasts under the CaMV promoter in earlier work (Mizutani et al., 2006; Choi and Roberts, 2007). Subsequent analysis of NIP2;1-GFP localization in transgenic Arabidopsis plant roots driven by a heterologous NIP5;1 promoter under aerobic conditions, also showed dual localization (Wang et al., 2017).

The trafficking of plant aquaporins to various target membranes through endocytic and redistribution pathways is regulated based on metabolic need and stress physiology [reviewed by (Chevalier and Chaumont, 2015; Takano et al., 2017)]. For example, PIP2;1 aquaporins are dynamically cycled between the internal membranes and the plasma membrane (Li et al., 2011), with regulation via phosphorylation (Boursiac et al., 2008; Prak et al., 2008) or other factors (Santoni, 2017; Takano et al., 2017) leading to preferential surface expression or internalization, which regulates the hydraulic conductivity of the cell. The dual localization of NIP2;1 could reflect a similar dynamic distribution and trafficking between internal membranes and the cell surface to regulate lactate efflux. In the case of some plant (Prak et al., 2008) as well as mammalian (Noda and Sasaki, 2006) aquaporins, preferential trafficking to the plasma membrane is controlled by the phosphorylation of serine within the cytosolic carboxyl terminal domain. Proteins of the NIP I subgroup are phosphorylated on a homologous serine within the carboxyl terminal domain (Wallace et al., 2006; Santoni, 2017), which is catalyzed by CDPK/CPK kinases (Weaver et al., 1991). This phosphorylation motif is conserved in NIP2;1 (Ser 278). In addition, phosphoproteomic analysis reveals that NIP2;1 is also phosphorylated in the N-terminal domain at Ser 5 by an unidentified protein kinase (Vialaret et al., 2014). Whether phosphorylation, or other regulatory factors, control trafficking or distribution of NIP2;1 in response to hypoxia or recovery signals to regulate lactic efflux or uptake remains to be investigated.

### Lactic acid and ethanolic fermentation pathways

Ethanolic fermentation through the PDC-catalyzed decarboxylation of pyruvate followed by subsequent production of ethanol from acetaldehyde via ADH is proposed to be the major anaerobic catabolism pathway (Gibbs and Greenway, 2003). However, lactic acid fermentation is also carried out in most plant species (Gibbs and Greenway, 2003), and in many cases may precede, and regulate the transition to ethanolic fermentation. The reason for initial reliance on lactic acid fermentation during hypoxia prior to a shift to ethanolic fermentation is not clear (Gibbs and Greenway, 2003). However, this pathway, unlike ethanolic fermentation, could allow recovery of the fermentation end product. Additionally, lactate production via LDH occurs under aerobic conditions in response to other abiotic and biotic stresses that could affect energy metabolism (Dolferus et al., 2008; Maurino and Engqvist, 2015). In animal systems, the physiological role of lactate transcends serving as an end product for anaerobic glucose metabolism, and its larger role as a metabolic regulator has emerged, including G-protein signalling as well as transcriptional regulation through histone modification (Latham et al., 2012; Sun et al., 2017; Zhang et al., 2019).

The possibility of lactate as a signal in plant systems remains largely unexplored. Nevertheless, there is evidence that the balance between the two fermentation pathways is regulated. For example, in classical studies of maize root tips, plants subjected to hypoxia stress initially engage in lactic acid fermentation followed by a switch to primarily ethanolic fermentation that is proposed to be driven by cellular acidification by lactate accumulation or other means that results in subsequent pH-dependent activation of PDC [the “pH stat” model (Davies et al., 1974; Roberts et al., 1984)]. In the case of Arabidopsis, overexpression of *LDH* results in an increase in PDC activity and media efflux of lactate (Dolferus et al., 2008), suggesting that increased lactic acid fermentation triggers the expression of ethanolic fermentation enzymes. Conversely, *adh1* null plants induce higher levels of lactic acid production to apparently compensate for reduced flux through the ethanolic fermentation pathway (Ismond et al., 2003). In *nip2;1* mutants, the accumulation of higher tissue lactate may produce an effect similar to *LDH1* overexpression. Higher transcript levels encoding enzymes within alternative pathways of the metabolism of pyruvate (e.g., *PDC1, ADH1* and *AlaAT1*) may be an adaptive response to the accumulation of lactic acid in *nip2;1* roots.

The reason for the selective reduction of *GOX3* inhibition in hypoxic *nip2;1* roots is less clear. As pointed out by Engqvist et al. (2015), the proposed role of this enzyme is to convert lactate back to pyruvate within the peroxisome which would serve to reduce lactic acid levels within the cell. Notably, however, this conversion occurs with the production of a reactive oxygen end product (hydrogen peroxide). ROS production is a major contributor to reoxygenation stress and is associated with poor tolerance to hypoxia and recovery (Yeung et al., 2019). If cytosolic lactate levels are elevated in *nip2;1* mutants, GOX3 (which uses oxygen as a co-substrate) could trigger greater ROS production upon reoxygenation.

### Aquaporins as lactic acid channels in other plant and microbial systems

In addition to Arabidopsis NIP2;1, select aquaporins with lactic acid permeability and efflux function have been described in other systems. For example, the Lactobacillales, which produce large quantities of lactic acid through fermentation, possess isoforms of the glycerol facilitator encoded by the *GlpF1* and *GlpF4* that facilitate lactic acid efflux (Bienert et al., 2013). The human trematode pathogen, *Schistosoma mansoni*, which performs lactic acid fermentation during the pathogenic part of its life cycle, possesses a lactic acid permeable plasma membrane aquaporin SmAQP that is proposed to release this end product (Faghiri et al., 2010). More pertinent to the present study, recent work (Mateluna et al., 2018) has identified other NIP I proteins, PruavNIP1;1 and PrucxmNIP1;1, that are induced during low oxygen stress in hypoxia-tolerant *Prunus* root stocks, and which are proposed to be lactic acid permeable proteins based on yeast lactate auxotroph assays. These observations suggest that a subset of the NIP I family have evolved to functionally flux lactic acid. Since NIP2;1 retains the ar/R pore constriction properties of other lactic acid-impermeable NIP aquaglyceroporins, the structural alterations within the pore that confer the altered selectivity for lactic acid and the exclusion of other substrates include additional pore determinants besides the canonical selectivity filter that remain to be elucidated. Finally, whether NIP proteins are part of a larger network of transporters that coordinate directional lactate efflux and movement within root tissues, similar to NIP and BOR proteins in boric acid homeostasis (Takano et al., 2008), remains unknown.

## Materials and Methods

### Plant growth conditions and stress treatments

*Arabidopsis thaliana* ecotype Columbia-0 was used in all experiments. Seeds were sterilized and stratified at 4 °C for 2 days, and were germinated as in (Choi and Roberts, 2007). Seedlings were grown vertically on in Murashige-Skoog (MS) media supplemented with 1% (w/v) sucrose and 0.8% (w/v) Phyto-agar (plantMedia) with a long day (LD) cycle of 16 hours of light (100 μmol m^-2^ s^-1^) and 8 hours of dark (LD conditions). Hypoxia treatment was administered at the end of the light cycle by the argon-treatment protocol described by (Lokdarshi et al., 2016). For normoxic controls, seedlings were treated simultaneously under identical conditions except in the presence of air instead of argon gas. Phenotype analysis for hypoxia survival was conducted as described by Lokdarshi et al. (2016) with modifications. Briefly, seven days old seedlings were administered 8 hours of argon gas-induced hypoxia or air, and were returned to LD growth conditions. The survival frequency (seedling chlorosis) was scored three days after stress treatment.

### Photosynthetic efficiency measurement

The maximum quantum yield of photosystem II [QY_max_= F_v_ /F_m_] was measured with a FluorCam 800MF instrument (Photon Systems Instruments) by the general method of (Murchie and Lawson, 2013). Seven-day old seedlings were administered 8 hr hypoxia, and QY_max_ was measured at different recovery time points upon return to LD conditions. For the first time point (time=0), seedlings were removed from hypoxia and were subjected to a saturating pulse of 1800 µEin m^-2^s^-1^ for 0.8 sec (F_m_). Variable fluorescence (F_v_) was calculated as the difference between F_o_ and F_m_ to calculate the maximum quantum yield [F_v_/F_m_]. For subsequent measurements, seedlings were dark adapted for 2 min (F_0_) prior to application of the saturating pulse and conducting measurements.

### T-DNA Insertion Mutant *nip2;1* and complementation line

A sequence tagged T-DNA insertion line within the *NIP2;1* gene (WiscDsLox233237_22k) from the WiscDs-Lox T-DNA collection (Woody et al., 2007) was obtained from Arabidopsis Biological Resource Center at the Ohio State University. The mutant, herein named *nip2;1*, was selected on MS media supplemented with 15 μg/mL Basta. The genotype of mutant plants was assessed by a PCR-based genotyping protocol as described at (http://signal.salk.edu/tdnaprimers.2.html). For this purpose, genomic DNA was extracted from 2-wk old seedlings using the *Wizard* Genomic DNA purification kit (Promega) and was subjected to PCR analysis using two *AtNIP2;1* gene specific primers and the left border T-DNA primer (all primers used in this study are described in Supplemental Table 1). The precise site of T-DNA insertion was verified by a cloning of the PCR product into the pCR2.1-TOPO vector (Invitrogen) followed by automated DNA sequencing. All sequencing was performed at the University of Tennessee Molecular Biology Resource Facility. T4 homozygous mutant seedlings were used for all phenotyping and other analyses in this study.

For complementation of *nip2;1*, as well as to provide a mechanism for localization of a native protein construct by fluorescence microscopy, transgenic lines containing a transgene consisting of a *NIP2;1-GFP* translational fusion under the control of the *NIP2;1* promoter were generated in the *nip2;1* background. The promoter region of the *NIP2;1* gene (from the translational start site to a site 2000 bp upstream) was generated by PCR of Arabidopsis genomic DNA with gene specific primers with added *Kpn*I and *AatII* restriction sites (Supplemental Table 1) to facilitate insertion into the binary vector pKGW_RedRoot_OCSA (Niyikiza et al., 2020). The modified destination vector was named pKGW_OCSA_*NIP2;1Pro*. The *NIP2;1* coding sequence (CDS) was amplified from the cDNA prepared from 4 hr hypoxic Arabidopsis seedlings with gene-specific primers with *Xba*I and *EcoR*I sites (Supplemental Table 1) to facilitate cloning into the Gateway entry vector CD3-1822 (Wang et al., 2013) to generate a construct of encoding NIP2;1 as an in-frame carboxyl-terminal fusion with GFP separated by a 3X Gly linker. The *NIP2;1-GFP* construct was then recombined into the pKGW_OCSA_*NIP2;1Pro* vector by using a gateway LR reaction (Invitrogen) to generate the final binary vector with *NIP2;1*_*pro*_::*NIP2;1*-*GFP*. The constructs were sequenced and verified by using Snap Gene 4.2.11 software. *Agrobacterium tumefaciens* GV3101 (Koncz and Schell, 1986) was transformed with the final construct by electroporation with a Bio-Rad Gene Pulser Xcell Electroporation system. Colonies carrying the correct construct were verified by PCR, and were used to transform Arabidopsis *nip2;1* plants by using floral dip method (Clough and Bent, 1998).Transgenic lines were selected on MS media supplemented with 25 μg/ml kanamycin, and were confirmed by PCR-based genotyping with transgene specific primers (Supplemental Table 1). Twelve transgenic lines were generated and the three with the highest *NIP2;1* transgene abundance were selected for further analysis. T_2_ generation complementation lines were used for the studies.

### RNA Purification and Quantitative PCR

Total RNA was isolated from plant tissues by grinding on liquid nitrogen and extracted by PureLink Plant RNA Reagent (Invitrogen). RNA isolation and DNase treatment was carried out by Direct-zol RNA MiniPrep Plus Kit using the manufacturer’s protocol (Zymo Research). Q-PCR was performed on a Bio-Rad IQ5 real-time PCR detection system by using iTaq Universal SYBR Green One-Step RT-qPCR kit (Bio-Rad Laboratories) according to the manufacturer’s instructions. All gene specific primers used in this study are summarized in Supplementary Table 1. Quantitative expression analysis was calculated by the comparative threshold cycle (Ct) method (Pfaffl, 2001). Either *UBQ10* or *ACTIN2* transcripts were used as the reference C_t_ for calculating ΔC_t_ for target gene expression, and standardization to a calibrator transcript (ΔΔC_t_) was done as described in (Choi and Roberts, 2007). The specific calibrators described in the figure legends.

### Histochemical and microscopy techniques

GUS staining and clearing of *Arabidopsis thaliana* lines with the *NIP2;1*_*pro*_:*GUS* reporter transgene was done by the protocol of (Choi and Roberts 2007), and stained tissues were imaged with a LEICA MZ16FA microscope (Leica Microsystems.). For analysis of *NIP2;1* promoter activity in root cross-sections, GUS-stained roots were dehydrated in ethanol and embedded in Technovit 7100 resin by the manufacturer’s (Kulzer GmbH) protocol. Cross sections (2.5 µm thickness) were generated from the mature differentiated region of the primary root with a Reichert OMV3 microtome equipped with a glass knife, and were mounted in 50% (w/v) glycerol. Cross-sections were imaged with a Nikon ECLIPSE E600 microscope equipped with Micropublisher 3.3 and QCapture 2.60 software (QImaging corporation).

Epifluorescence imaging of NIP2;1-GFP plants were captured with an Axiovert 200M microscope (Zeiss) equipped with filters for GFP fluorescence (Zeiss; filter set 38 HE) and a digital camera (Hamamatsu Orca-ER) controlled by the Openlab software (Improvision). Subcellular localization analysis under hypoxia and reoxygenation was done by confocal microscopy. Hypoxia treatment was done with the argon protocol described above, and seedlings were removed from the plate and were incubated in 4 uM FM4-64 under hypoxic conditions (Invitrogen) in darkness for 10 min. For reoxygenation, seedlings were returned to aerobic conditions under light for 1 hr before staining and visualization. Imaging was performed with a Leica SP8 white laser confocal microscope system at the Advanced Microscopy and Imaging Center at The University of Tennessee, Knoxville. The 488-nm excitation filter was used, and the emission filter for detection was set to 495 to 550 for GFP and 580 to 650 nm for FM4-64. Confocal micrographs were captured with the Leica LASX software and uncompressed images were exported and analyzed in ImageJ version 1.53a (Schneider et al., 2012) to adjust the brightness and contrast of images, and to generate merged images.

### Immunochemical Techniques

Anti-NIP2;1 antisera were produced against a synthetic peptide (GenScript) corresponding to the C-terminal sequence of NIP2;1 (CHKMLPSIQNAEPEFSKTGSSHKRV) following the immunization protocol of (Guenther et al., 2003) with the exception that Titermax-Gold was substituted for Freund’s adjuvants. Antibodies were affinity purified on peptide resins as described in (Guenther et al., 2003).

For analysis of NIP2;1 protein in wild type and *nip2;1* mutant after hypoxia treatment, Arabidopsis root tissues from seedlings treated with 6 hr hypoxia or normoxic controls were extracted, a membrane microsomal fraction was prepared as described by (Ishikawa et al., 2005). Protein concentrations were determined by using the BCA assay (Pierce Biochemical). The SDS-gel electrophoresis and Western blot analysis were performed using 10 μg protein from membrane microsomal fractions as previously described (Guenther et al., 2003). For the analysis of NIP2;1-GFP expression in complementation lines, hypoxia and reoxygenation treatments were conducted as described above, and samples were directly extracted into SDS-PAGE sample buffer (Laemmli, 1970) for Western blot analysis. Rabbit anti-GFP polyclonal antibodies (Abcam) were used for the detection of NIP2;1-GFP.

### Media Acidification and L-Lactate Measurements

Media acidification assays with the pH-sensitive indicator bromocresol purple was done by the method of (Silva et al., 2018). Seven day wild type and *nip2;1* seedlings were transferred from MS media to 1.5% agarose plates containing 60 mg L^-1^ Bromocresol Purple (Acros Organics) and 1mM CaSO_4_ in sterile distilled water with the pH adjusted to 5.7 using KOH. The bromocresol purple plates with the seedlings were subjected to anaerobic condition as described above for 8 hrs before the images were captured with a DSLR camera (Canon Rebel XS).

To quantify the L-lactate efflux from roots to the external media, seven-day old wild type and *nip2;*1 7-d-old seedlings were transferred to Eppendorf tubes, with roots submerged in 1mL of HPLC water, and were subjected to argon gas treatment as described above. At the selected time points the plant material was removed and the L-lactate levels were quantified enzymatically (LDH) as described by (Marbach and Weil, 1967). To determine L-lactate concentration within Arabidopsis roots, hypoxia treatment of seedlings was carried out, roots were dissected, and tissues were snap frozen in liquid nitrogen. The frozen tissue was ground in a mortar with a pestle and was extracted in two volumes of 1N perchloric acid and was neutralized with potassium carbonate on ice prior to assay by the LDH method of (Bergmeyer and Bernt, 1974). NADH production was assayed by the change in absorbance at 340 nm, with background corrected from duplicate samples treated identically except for no added LDH enzyme.

### Gene Accession Information

Accession number of the genes used in this study are AT2g34390 (*NIP2;1*), AT4g33070 (*PDC1*), AT4g18360 (*GOX3*), AT1g17290 (*AlaAT1*), AT1g77120 (*ADH1*), AT3g18780 (*ACTIN 2*), AT4g05320 (*UBIQUITIN 10*), AT4g17260 (*LDH1*).

## Supplemental data

**Supplemental Figure 1** Cellular localization of *AtNIP2;1* promoter activity in the roots under normoxic growth conditions.

**Supplemental Figure 2** Complementation of *nip2;1* T-DNA mutant with a *NIP2;1pro::NIP2;1-GFP* construct.

**Supplemental Figure 3** Subcellular localization of NIP2;1-GFP in the roots of hypoxia challenged 7-d-old *NIP2;1-GFP* complementation lines.

**Supplemental Figure 4** Comparison of hypoxia-induced expression of *NIP2;1* and lactate metabolizing enzyme transcripts *LDH* and *GOX3*

**Supplemental Table 1** Oligonucleotide primers

## Acknowledgements

We appreciate the assistance and valuable contributions of the following undergraduate students of the BCMB Department at the University of Tennessee, Knoxville: Samantha McIntire for helping in phenotyping experiment, and Shivam Ishanpara for helping us in growing and genotyping transgenic plants and preparing media for the experiments; and especially Clayton Nunn who participated in initial localization experiments. We are thankful to Tessa Burch-Smith and John Dunlap for assisting us in microscopy in initial studies, and Andreas Nebenfuehr for allowing us to use his laboratory’s dissecting and epifluorescence microscopes.

